# Regional Glymphatic Dysfunction is linked to Spinocerebellar Ataxia Type 3 pathophysiology

**DOI:** 10.1101/2024.04.16.589724

**Authors:** Lin Hua, Manxi Xu, Linwei Zhang, Fei Gao, Xinglin Zeng, Aocai Yang, Jixin Luan, Amir Shmuel, Guolin Ma, Zhen Yuan

**Affiliations:** Faculty of Health Sciences, University of Macau, Macau SAR, China; Centre for Cognitive and Brain Sciences, University of Macau, Macau SAR, China; Department of Radiology, China-Japan Friendship Hospital, Beijing, China; Peking University China-Japan Friendship School of Clinical Medicine, Beijing, China; Department of Neurology, China-Japan Friendship Hospital, Beijing, China; Institute of Modern Languages and Linguistics, Fudan University, Shanghai, China; China-Japan Friendship Hospital (Institute of Clinical Medical Sciences), Chinese Academy of Medical Sciences & Peking Union Medical College, Beijing, China; McConnell Brain Imaging Centre, Montreal Neurological Institute, McGill University, Montreal, QC, Canada; Departments of Neurology and Neurosurgery, Physiology, and Biomedical Engineering, McGill University, Montreal, QC, Canada

**Keywords:** cerebrospinal fluid flow, resting-state fMRI signal, choroid plexus, DTI-ALPS, glymphatic system, spinocerebellar ataxia type 3

## Abstract

Spinocerebellar ataxia type 3 (SCA3) involves neuroinflammation and imbalance between production and clearance of proteins which affects the glymphatic system, the lymphatic-like, fluid-transport system in the brain. However, it is unclear whether SCA3 is related to impairments in glymphatic function. Using multimodal imaging data, 34 SCA3 patients and 36 age-, sex- and educational matched healthy controls (HCs) were compared using multiple glymphatic measurements, including choroid plexus (CP) and cerebrospinal fluid (CSF) volume, diffusion tensor imaging along the perivascular (DTI-ALPS) index, and coupling relationship between blood-oxygen-level-dependent signals and CSF flow (BOLD-CSF coupling). Then, we evaluated regional glymphatic function by dividing DTI-ALPS and BOLD-CSF coupling into anterior, middle, posterior, and cerebellum regions, thereby identifying the spatial variation of glymphatic function in the two groups. We demonstrated that compared with HCs, larger CP and CSF volumes were found in SCA3 patients. More importantly, for DTI-ALPS index and BOLD-CSF coupling, these surrogate markers for glymphatic clearance were weaker in SCA3 patients. Furthermore, altered regional glymphatic functions were most prominent in midbrain, cerebellum and middle regions. Crucially, the altered midbrain, cerebellum, middle and global glymphatic functions were accompanied by the severity of ataxia and other SCA3 symptoms. Similar to other neurodegenerative disorders, the association between multiple glymphatic indexes and SCA3 symptoms suggested that waste clearance is disrupted in SCA3 patients, which shed light on the pathogenesis of this disease from a glymphatic lens. Our findings highlighted the dysregulated glymphatic function as a novel diagnostic marker for SCA3.

## Introduction

Spinocerebellar ataxia type 3 or Machado-Joseph disease (SCA3/MJD) is the most common autosomal dominant cerebellar ataxia in neurodegenerative diseases. Individual with SCA3 typically experience motor and non-motor symptoms, including pyramidal signs, extrapyramidal features, cognitive decline, and rapid eye movement behavior disorder (RBD)^[1–4]^. The pathogenesis of SCA3 is thought to be the protein misfolding, wherein the expansion of CAG repeats within the coding region of the ATXN3 gene, sited on chromosome 14, and engenders abnormal polyglutamine proteins^[5, 6]^. This dysregulation, distorting the equilibrium between protein production and clearance, impairs cell function and precipitates neuron death. Sustaining homeostasis in the brain environment is crucial for regulating and supporting the mechanisms to clear abnormal proteins^[7, 8]^. This perturbation affects the cell environment, neuroinflammation, the evolution of complex pathology and disease progression. Additionally, what play key roles in the SCA3 mechanism are increased inflammatory responses, accumulation of metabolic waste due to mitochondrial dysfunction, and imbalances in regulating cell stress responses^[9]^. Given the role of these pathological processes, the brain’s waste clearance, particularly by the glymphatic system, is critical for maintaining the health of the nervous system. However, research on assessing glymphatic system in the context of SCA3 is limited. It is thus imperative to understand the pathophysiology of SCA3. To address this issue, this study attempts to investigate the functional status of the glymphatic system in SCA3 patients, and its impact on disease development and progression.

The glymphatic system, as the central nervous system’s internal “waste clearance” mechanism, facilitates the exchange between CSF and interstitial fluid (ISF) and thus effectively removes metabolic waste and abnormal proteins^[7, 8]^. CSF is introduced into the brain tissue via periarterial pathways while ISF clears through perivenous spaces, using astrocytic Aquaporin 4 (AQP4) water channels, which enables dynamic fluid exchange between brain tissue and perivascular spaces (PVS)^[10–12]^. This mechanism not only supports the distribution of nutrients and the transport of medications but also plays a significant role in the pathogenesis of neurodegenerative diseases, such as Alzheimer’s disease (AD) ^[13, 14]^ and Parkinson’s disease (PD) ^[15, 16]^. Neuropathological and MRI studies have revealed widespread abnormal protein accumulation and neuronal loss in SCA3. Specifically, the nuclear accumulation of polyglutamine is distributed in the cerebral cortex, cerebellum cortex, basal ganglia, and thalamus^[17, 18]^. The rise in serum neurofilament light chain (NfL) levels, indicative of neuronal damage and axonal degeneration, directly mirrors the extent of neuronal injury and the clinical severity in SCA3 patients ^[19, 20]^. Furthermore, CSF t-tau and p-tau^181^ have been reported to be elevated in the early stages and subsequently decreased over time in SCA3 patients^[21]^. A compromised glymphatic system may result in a diminished clearance of abnormal proteins and neuro-damage markers like NfL, potentially worsening the neuronal damage triggered by neurodegenerative diseases. Additionally, it is noteworthy that the activities of the glymphatic system increase during sleep, a critical period for the clearance of waste from the brain^[22]^. However, SCA3 patients often experience difficulties in falling asleep and frequent awakenings^[3, 4]^, which may disrupt the normal glymphatic function and further leads to reduced efficiency in removing waste from the brain. Collectively, these findings indicated that impaired glymphatic function and neuroinflammation may be foundational to the progression of SCA3-related diseases, illuminating the pathogenic mechanisms of this disease through an exploration of glymphatic function.

Nevertheless, it has been challenging for neuroimaging studies to examine glymphatic impairments in SCA3 due to the lack of tools for accessing glymphatic clearance-related activities. A pioneering study revealed a strong coupling between global low-frequency (<0.1 Hz) resting-state fMRI (rs-fMRI) BOLD signals and CSF dynamics in the ventricles during sleep^[22]^, which may contribute to metabolite clearance^[23]^. Moreover, alterations in BOLD-CSF coupling in the wakeful resting state have been found in patients with AD^[24, 25]^, PD^[26, 27]^, and Frontotemporal Dementia (FTD)^[28]^. Furthermore, CP is the primary locus of CSF secretion and serves as the gateway for the trafficking of the inflammatory cells during central nervous system (CNS) injuries and infections^[29, 30]^. Recent studies pointed that functional dysregulation of the CP-CSF reflected the common underlying mechanism in the pathophysiology of neurodegenerative diseases^[30]^. Dysfunction of the CP and CSF flow could precede the neurological symptoms and affect glymphatic function in neurodegenerative diseases^[28, 31, 32]^. Meanwhile, DTI-ALPS index is also a recently established non-invasive method to quantify the dynamics of subcortical glymphatic flow, especially water movement along the periventricular pathway, which is associated with the driving force of glymphatic function^[33]^. Reduced DTI-ALPS indexes in patients with AD^[33, 34]^ and PD^[35, 36]^ were associated with increased disease severity and attributed to reduced glymphatic function. These evidence converged to support the notion that reduced glymphatic clearance is implicated in neurodegenerative diseases^[37]^. Therefore, we speculated that SCA3 might also involve changes in CP and CSF morphology, BOLD-CSF coupling, and DTI-ALPS, which quantifies glymphatic dysfunction and neuroinflammation.

Against this background, the current study capitalized on multimodal imaging data (structural MRI, rs-fMRI, and DTI data) and aimed to comprehensively understanding the glymphatic dysfunction in SCA3. The alterations in CP and CSF morphology, the coupling between the BOLD signal and ventricle CSF flow, and DTI-ALPS index were evaluated. Additionally, to investigate the spatial pattern changes of glymphatic system in SCA3 patients, glymphatic measurements was separated into regional BOLD-CSF coupling and regional DTI-ALPS from anterior to posterior and further to cerebellum regions. Finally, the association between glymphatic measurements and neuropsychological assessments in SCA3 patients would be calculated.

## Methods

### Subjects

The present cohort consists of 70 subjects (44.21±12.64 years; 28 females) categorized into HCs (*N* = 36) and SCA3 patients (*N* = 34). The patients were recruited from the outpatient clinic of the China-Japan Friendship Hospital (see Supporting information for detailed subjects’ information) while the HCs from the local communities. All subjects underwent clinical assessments, T1-weighted structural MRI, rs-fMRI and DTI scans (Figure 1A). The SCA3 patients were screened by family history, ataxia assessments including the Scale for the Assessment and Rating of Ataxia (SARA) and the International Cooperative Ataxia Rating Scale (ICARS)^[38]^, genetic and molecular tests, and routine brain MRI findings to validate diagnoses by expert neurologists. The HCs were matched for age, sex, and educational level with SCA3 patients, free of any neuropsychiatric or neurodegenerative diseases. In addition to ataxia assessments, SCA3 patients were performed extraordinary psychiatric assessments using Self-Rating Anxiety Scale (SAS)^[39]^ and Self-Rating Depression Scale (SDS)^[40]^. Additionally, to evaluate the SCA3 patients’ sleep quality in, Self-Rating Scale of Sleep (SRSS) was also involved^[41]^. Finally, after controlling the imaging quality, 36 HCs and 34 SCA3 patients with structural MRI, 35 HCs and 27 SCA3 patients with rs-fMRI data and 35 HCs and 28 SCA3 patients with DTI data were selected for further CP and CSF volumes, BOLD-CSF coupling and DTI-ALPS analysis, respectively (see Supporting information for Quality control).

**Figure 1.**
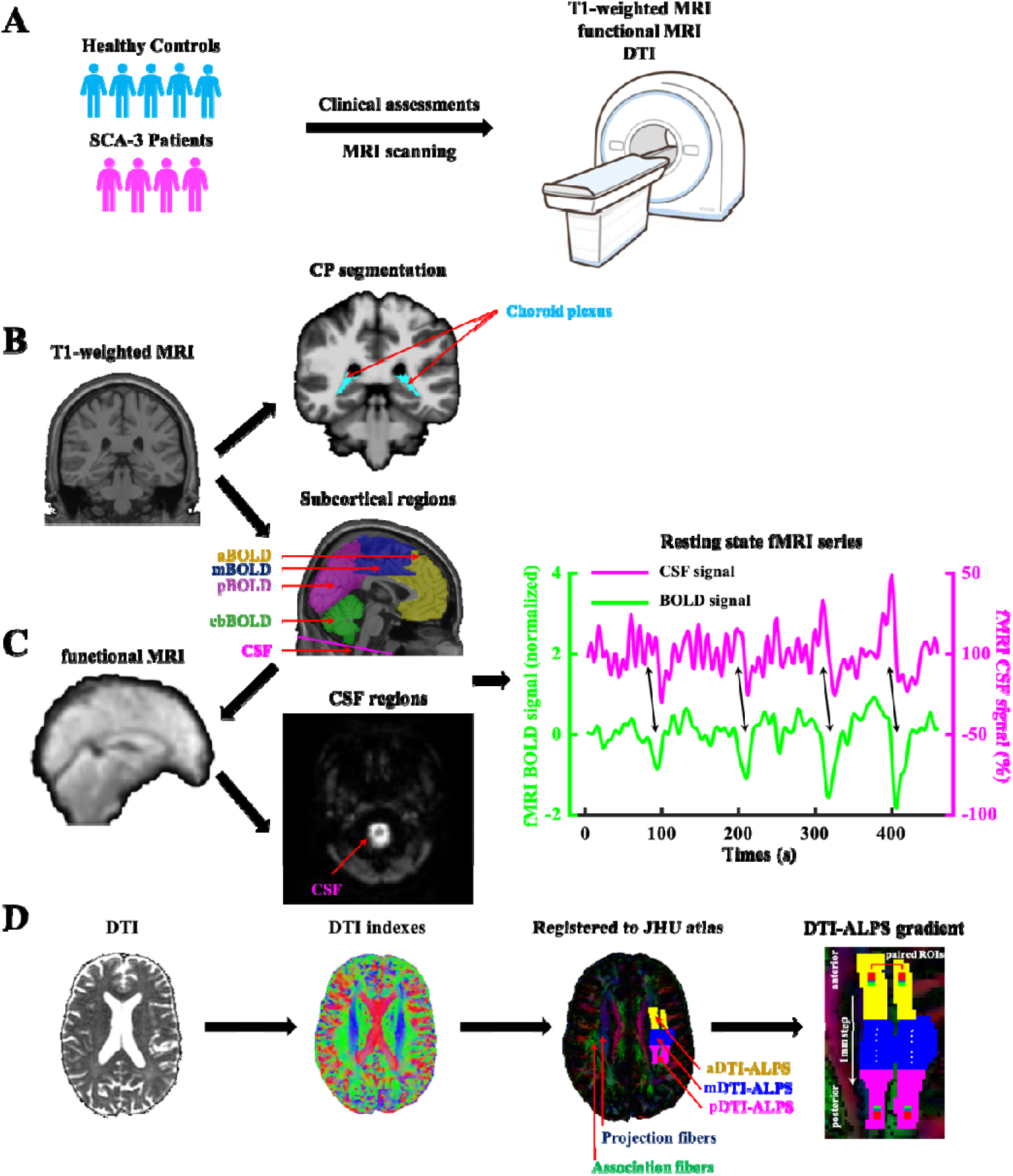
Whole procedures of characterizing multiple glymphatic measurements. (A) The present cohort of 70 subjects were categorized in HC (*N* = 36) and SCA3 (*N* = 34) groups, and underwent neuropsychological assessments, T1-weighted structural MRI, rs-fMRI and DTI scans. (B) Choroid plexus within the lateral ventricles and cerebrospinal fluid were automatically segmented and manually corrected in each subject’s structural MRI. (C) The global BOLD signal was extracted from gray matter regions (including the cerebellum), and then further separated into anterior BOLD, middle BOLD, posterior BOLD, and cerebellum BOLD regions. The CSF signal was extracted from CSF regions at the bottom slice of the rs-fMRI image and referenced to the structural MRI image. CSF inflow effect was detected to identify the boundaries of CSF regions at the bright areas of the rs-fMRI image. Representative time series of normalized cortical gray-matter BOLD signal and CSF signal changes were showed by corresponding amplitude fluctuations. (D) DTI images were preprocessed and calculated to generate the FA map. Then, the FA map was registered to the JHU atlas and segmented into anterior DTI-ALPS, middle DTI-ALPS, and posterior DTI-ALPS regions according to the atlas labels (anterior, superior, posterior corona radiata, and superior longitudinal fasciculus). The glymphatic gradient was calculated by the DTI-ALPS from 46 pairs of 3 × 3 × 3 mm^3^ ROIs with 1 mm intervals from anterior to posterior in the projection and association fibers.

### Image acquisition and preprocessing

All MRI data (structural MRI, rs-fMRI, and DTI data) was collected in the 3.0 Tesla MR scanner (General Electric, Discovery MR750, Milwaukee, WI, United States). Each imaging session included a three-dimensional fast spoiled gradient-echo sequence (3D FSPGR) with the following parameters: repetition time (TR) = 6.7 ms, echo time (TE) = 2.9 ms, flip angle = 12°, slice thickness = 1 mm, field of view (FOV) = 256 × 256 mm^2^, matrix = 256 × 256, voxel size = 1 × 1 × 1 mm^3^. For rs-fMRI acquisition, 240 fMRI volumes were captured with an echo-planar image (EPI) sequence (TR = 2000 ms, TE = 30 ms, filp angle = 90°, slice thickness = 3.5 mm, FOV = 224 × 224 mm^2^, matrix = 64 × 64, voxel size = 3.5 × 3.5 × 4.2 mm^3^). The DTI images with 8 b-value of 0 s/mm^2^ and 64 b-value of 1000 s/mm^2^ were acquired using a diffusion weighted spin-echo EPI (TR = 8028 ms, TE = 81.8 ms, flip angle = 90°, slice thickness = 2 mm, FOV = 240 × 240 mm^2^, matrix = 120 ×120, voxel size = 2 × 2 × 2 mm^3^). During the MRI scanning session, foam paddings were given inside the head coil to restrict potential head motions. Additional routine MR sequences, including axial T2-weighted imaging (T2WI), T2-FLAIR, and diffusion-weighted imaging (DWI), were performed to identify brain abnormalities in SCA3 patients.

The rs-fMRI data was preprocessed using the 1000 Functional Connectomes Project script (version 1.1-beta; https://www.nitrc.org/frs/?group_id=296), which underwent a pipeline similar to a previous study examining neurofluid coupling in neurodegeneration diseases^[24–28]^. Raw rs-fMRI images for each subject underwent preprocessing steps including slice timing, motion correction, skull stripping, spatial smoothing with a 4mm full-width half-maximum (FWHM) kernel, 0.01-0.1 Hz bandpass filtering, as well as linear and quadratic temporal trend removal. Subsequently, rs-fMRI images of the subjects were co-registered to their high-resolution T1-weighted structural MRI images and then to the 152-brain Montreal Neurological Institute (MNI-152) space. As our analysis focused on the coupling between CSF signal and global BOLD (gBOLD) signal, regression of global signal and CSF signal was omitted by our preprocessing pipeline. Additionally, under the same rationale regression of motion parameters was also skipped as it may associate with gBOLD signal^[42]^.

The DTI data was preprocessed using the FSL package (version 6.0.5; https://fsl.fmrib.ox.ac.uk/fsl/fslwiki). The general procedures for DTI data preprocessing included skull stripping, the correction for head motion, eddy current-induced distortions, EPI-induced susceptibility distortions, and bias field. The fractional anisotropy (FA) map of each subject was then registered to the FA map of the Johns Hopkins University atlas (JHU atlas) template^[43]^, and the transformation matrix obtained from the previous step of co-registration was applied to other diffusion metric maps to obtain all DTI-based maps in the space of the JHU atlas (spatial resolution, 1 × 1 × 1 mm^3^; Figure 1D).

### CP and CSF segmentation

The CP has been reported as the principal locus of CSF secretion, thus being considered as a potential imaging marker for indirectly evaluating CSF production and toxic clearance. Using the FreeSurfer cortical reconstruction pipeline (version 6.0.0; https://surfer.nmr.mgh.harvard.edu/fswiki), segmentation of the CP within the lateral ventricles was automatically accomplished from the T1-weighted structural MRI image (Figure 1B). Manual scrutiny and correction of CP segmentation was conducted by two neuroimaging researchers, ensuring its accuracy and reliability. These revised CP segmentations underwent further independent quality assessment and finalization by one neuroradiologist. Total intracranial volume (ICV), cortical gray matter (GM) volume, CSF volume, and CP volume were extracted from the whole brain segmentation. To reduce inter-subject variability, CP and CSF volumes were separately normalized as the ratio of CP and CSF volumes to ICV^[31]^.

### Quantification of the coupling between BOLD signal and CSF inflow

The gBOLD signals were obtained from the cortical GM regions of cerebrum and cerebellum, delineated by the automated anatomical labeling 2 (AAL2) atlas^[44]^ from preprocessed functional images in individual spaces (Figure 1C). To assess the regional distribution of glymphatic function, the AAL2 atlas was divided into the anterior, middle, and posterior parts based on the atlas labels of the frontal, temporoparietal, occipital, and cerebellum regions (Figure 1C). To ensure precise alignment between the cortical GM masks and the functional space, the cortical GM masks were transformed from the MNI-152 space to individual functional space for each subject using the inversed concatenated transformation matrix from the previous co-registration process (functional-structural-standard). Transformed cortical masks were visually inspected to assure accurate spatial registration with the preprocessed functional images. The CSF mask was placed in the bottom slices of the rs-fMRI image^[24–28]^ (Figure 1C). These bottom slices consistently encompassed the cerebellum of all subjects as confirmed by visual inspection, capturing the CSF through-slice inflow effect as previously described.

To quantify gBOLD-CSF coupling, the cross-correlation function was computed between the gBOLD signals and the CSF signals across different time lags ranging from -9s to 9s for each subject (Figure 1C). The negative correlation coefficients peaked at the lag of +3s. This negative peak signified the point at which the gBOLD and CSF signals exhibited their strongest inverse relationship, capturing the interplay between hemodynamic signal and CSF dynamics. It was therefore used to quantify the strength of the gBOLD-CSF coupling for each subject^[22]^. Moreover, we also calculated the cross-correlation function between the negative derivative of the BOLD signal and the CSF signal to ensure that the CSF signal matched the negative derivative of the BOLD oscillation when setting the negative value to zero^[22]^. Finally, the cross-correlation function between the CSF signal and the anterior, middle, posterior, cerebellum BOLD signals was computed to obtain the regional BOLD-CSF coupling (aBOLD-CSF, mBOLD-CSF, pBOLD-CSF, and cbBOLD-CSF couplings).

### Calculation of the DTI-ALPS index

The DTI-ALPS index, evaluating the water diffusivity along the PVS at the level of the lateral ventricle, is used to reflect the activity of the glymphatic system^[33–36]^. The direction of PVS at the lateral ventricle level is primarily along the x-axis, perpendicular to the ventricle wall. Furthermore, the projection fibers that pass along the lateral ventricular wall in the z-axis are adjacent to the association fibers that go through at the more lateral area in the y-axis. Therefore, the major distinction in water molecule behavior between diffusivity along the x-axis in both fibers (D_xx,proj_ and D_xx,assoc_) and diffusivity perpendicular to them (D_yy,proj_ and D_zz,assoc_) relates to the diffusivity of the PVS. The regions of interests (ROIs) in the current study were defined as 3-mm-diameter spherical areas within the projection (including the anterior, superior, and posterior corona radiata) and the association fiber (superior longitudinal fasciculus), localized at the level of the lateral ventricle (MNI coordinates z = 26) based on JHU atlas labels. Mean diffusion values of these ROIs were extracted to compute the global DTI-ALPS. Specifically, the DTI-ALPS index was calculated as follows^[33]^:

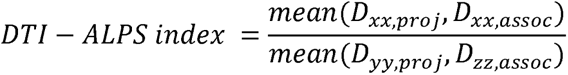

Subsequently, these ROIs were then subdivided into the anterior, middle, and posterior regions to obtain the anterior DTI-ALPS (aDTI-ALPS), middle DTI-ALPS (mDTI-ALPS), and posterior DTI-ALPS (pDTI-ALPS), respectively. Finally, DTI-ALPS gradient was calculated from the ROIs with 3 × 3 × 3 mm^3^ at 1 mm intervals to comprehensively understanding the glymphatic activity from anterior to posterior. A total of 46 pairs of ROIs were generated, and each pair of DTI-ALPS was calculated.

### Relationship between regional glymphatic measurements and neuropsychological assessments

The strength of gBOLD-CSF coupling was identified at the lag of +3s (the negative peak of the cross-correlation function). The gBOLD-CSF coupling strength at this lag was first correlated with age and compared between sexes. Then, confounding effects of age and sex were estimated by the linear mixed effect model and regressed out from further relationship analyses. Next, partial correlation analyses were conducted to evaluate the association within multiple regional glymphatic measurements in the whole cohort and in the two groups separately. Finally, the correlations between neuropsychological assessments, performed in SCA3 patients to assess ataxia, psychiatry as well as sleep symptoms, and CP ratio, CSF ratio, a/m/p/cb/gBOLD-CSF coupling strength and a/m/p/gDTI-ALPS strength after adjusting for age and sex.

### Statistical analysis

To ascertain the statistical significance of the a/m/p/cb/gBOLD-CSF correlations, permutation method was employed to create a null distribution for the a/m/p/cb/gBOLD-CSF correlations, respectively. Specifically, a/m/p/cb/gBOLD signals and CSF signals from different subject were randomly paired, and cross-correlation coefficients were recalculated. This process was repeated for 1000 times, generating a null distribution for the mean a/m/p/cb/gBOLD BOLD-CSF cross-correlation function. The *p* value was estimated by calculating the percentage of the correlation value of permutation data higher than the correlation value of real data at different lags.

Between-group comparisons were performed by two-sample t-test for continuous variables and χ^2^ test for categorical variables. Furthermore, Pearson’s correlation was used for the variables with normal distribution, while Spearman’s correlation was conducted to the non-normally distributed variables. Multiple comparisons were corrected by the false discovery rate (FDR) approach. Statistical significance was defined as *p* < 0.05.

## Results

### Demographic and clinical characteristics

A total of 70 subjects (44.21±12.64 years; 28 females) including 34 SCA3 patients (42.62±12.01 years; 10 females) was entered in the formal analyses (Table 1). 36 HCs (45.72±13.2 years; 18 females) were age- (*t*_(68)_ = 1.03, *p* = 0.308), sex- (*χ^2^*_(1)_ = 3.09, *p* = 0.079) and education- (*t*_(64)_ = 1.38, *p* = 0.171) matched with SCA3 patients. No significant differences were found in ICV (*t*_(68)_ = -0.57, *p* = 0.572) and GMV (*t*_(68)_ = 1.35, *p* = 0.183) between HCs and SCA3 patients. The mean distributions of disease durations and CAG repeat numbers in SCA3 patients were 9.04 ± 8.5 and 70.39 ± 3.67, respectively. Moreover, SCA3 patients underwent multiple neuropsychological assessments, including ataxia assessments of SARA (17.12 ± 9.2) and ICARS (44.74 ± 21.45), psychiatric assessments of SAS standard score (44.87 ± 12.45) and SAS standard score (47.1 ± 13.96), and sleep assessment of SRSS (26.32 ± 8.6).

**Table 1.**
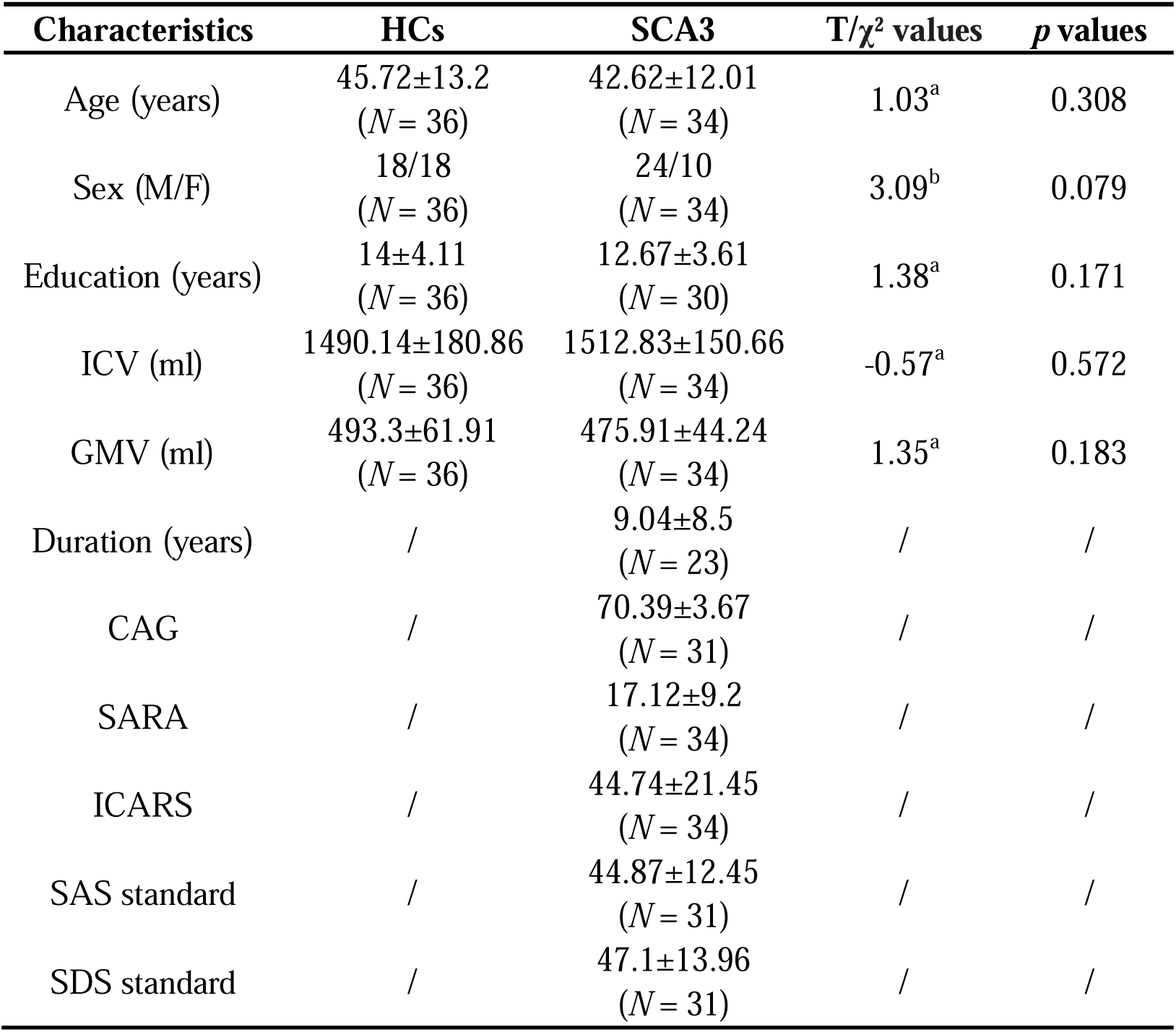
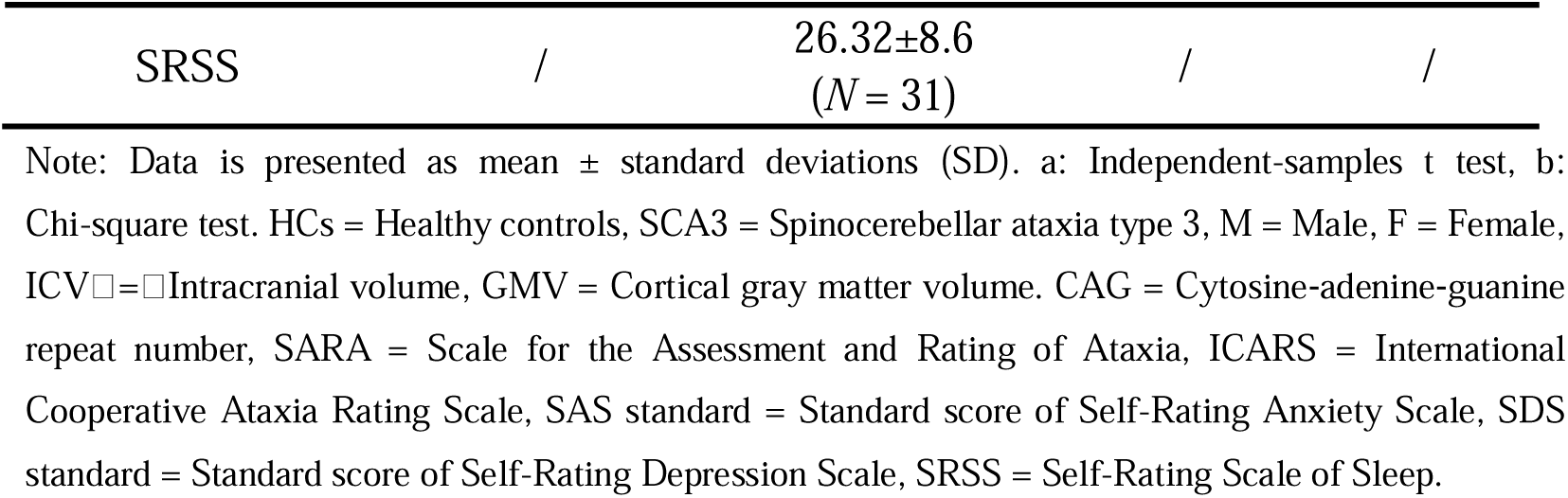
Demographic and clinical characteristics of HCs and SCA3 patients.

### The BOLD-CSF coupling alteration in SCA3

The gBOLD-CSF coupling was initially quantified by cross-correlation coefficient between gBOLD and CSF signals and averaged within HCs and SCA3 patients. Then, a/m/p/cbBOLD-CSF was obtained from the anterior, middle, posterior, and cerebellum regions of the BOLD signal coupled with the CSF signal, respectively. Consistent with previous report^[22, 24–28]^, the strength of gBOLD-CSF coupling for both groups (*r* = -0.29, *p* < 0.001 for HCs; *r* = -0.22, *p* < 0.001 for SCA3 patients; 1000 times permutation test) and a/m/p/cbBOLD-CSF coupling for the whole samples (*r* = -0.25, *p* < 0.001 for aBOLD-CSF coupling; *r* = -0.28, *p* < 0.001 for mBOLD-CSF coupling; *r* = -0.26, *p* < 0.001 for pBOLD-CSF coupling; *r* = -0.21, *p* < 0.001 for cbBOLD-CSF coupling; 1000 times permutation test) was identified with the negative peak of cross-correlation function at the lag of +3 (Figure 2A and 2B). At the individual level, the coupling strength did not correlate with head motion as quantified by mean frame-wise displacement (*r* = 0.12, *p* = 0.356; Figure S1). The peak of coupling strength and time lag was further confirmed by calculating the cross-correlation function between the negative derivative of the gBOLD signal and the CSF signal within HCs (Figure 2C) and SCA3 patients or between the negative derivative of the a/m/p/cbBOLD signal and the CSF signal across the entire sample (Figure 2D). Therefore, for further comparisons between two groups, the cross-correlation strength at +3s for each subject was used as a surrogate marker for glymphatic function.

**Figure 2.**
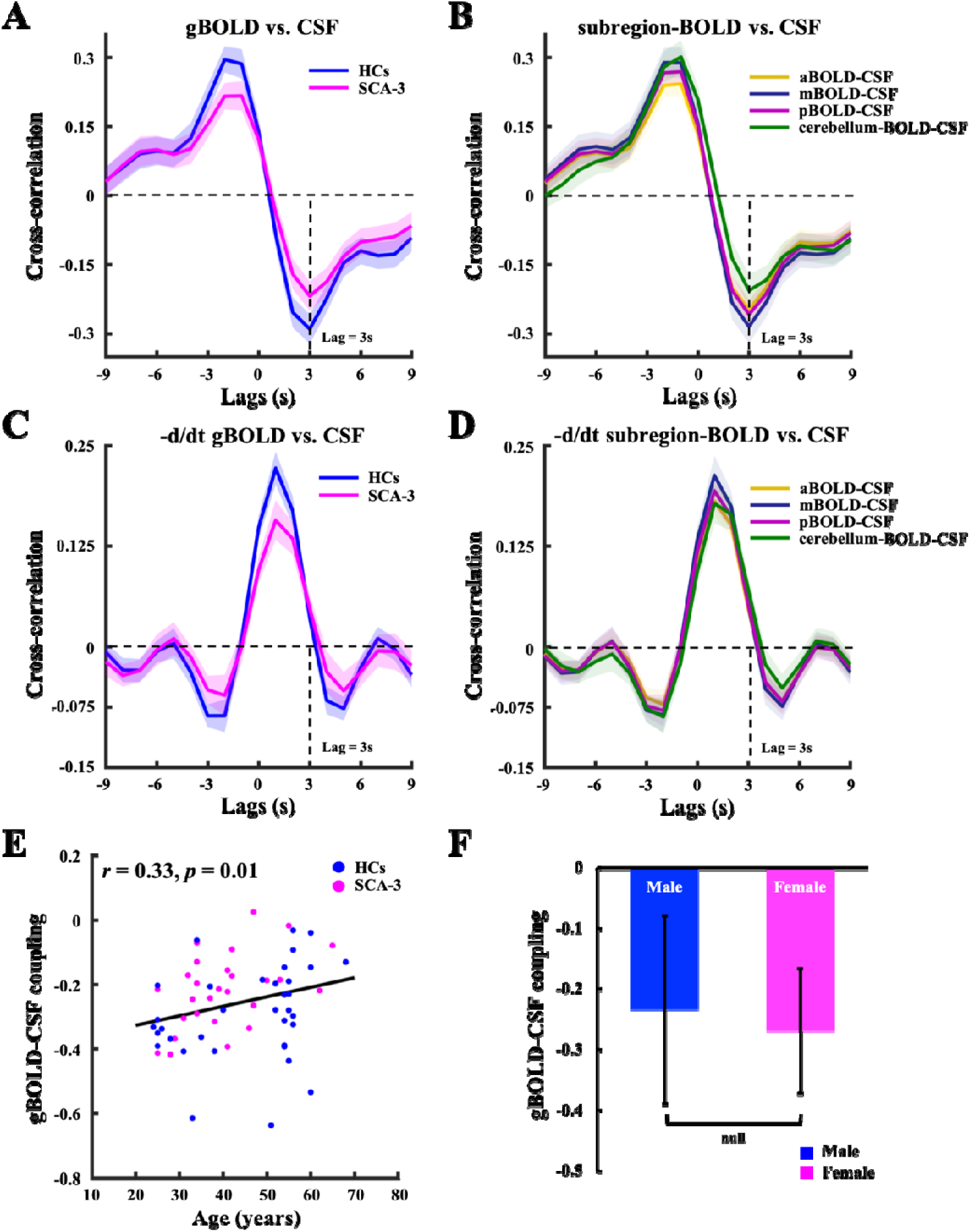
The BOLD-CSF coupling strength is associated with age, and differed between HC and SCA3 groups, as well as across brain regions. (A) The mean gBLOD-CSF cross-correlation function averaged within HCs (*N* = 35; blue line) and SCA3 patients (*N* = 27; magenta line). The vertical dashed line marks the +3s time lag (*r* = -0.29, *p* < 0.001 for HCs’ gBOLD-CSF coupling; *r* = -0.22, *p* < 0.001 for SCA3 patients’ gBOLD-CSF coupling; 1000 times permutation test; the negative peak of the mean cross-correlation). (B) The mean a/m/p/cbBLOD-CSF cross-correlation function averaged across all subjects (*N* = 62). The vertical dashed line marks the same +3s time lag as gBOLD-CSF coupling (*r* = -0.25, *p* < 0.001 for aBOLD-CSF coupling; *r* = -0.28, *p* < 0.001 for mBOLD-CSF coupling; *r* = -0.26, *p* < 0.001 for pBOLD-CSF coupling; *r* = -0.21, *p* < 0.001 for cbBOLD-CSF coupling; 1000 times permutation test). Dark yellow, navy blue, deep purple, and dark green indicate a/m/p/cbBOLD-CSF coupling, respectively. (C) and (D) Mean cross-correlation between the zero-threshold negative derivative of a/m/p/cb/gBOLD and CSF signals showed the strongest correlation at 3s (vertical dashed line). Shaded areas are 95% interval of the mean correlation coefficient across subjects. (E) The strength of gBOLD-CSF coupling, quantified as the cross-correlation at +3s, presented significantly positive correlation (Spearman’s correlation, *r* = 0.33, *p* = 0.01) with age across all subjects (*N* = 62). Blue and magenta dots indicate HCs (*N* = 35) and SCA3 patients (*N* = 27). (F) Male and female subjects showed no different gBOLD-CSF coupling in the whole cohort (*t*_(60)_ = 1.013, *p* = 0.315; Independent-samples t test). Blue and magenta bars indicate males (*N* = 36) and females (*N* = 26). Error bars represent the standard deviation (SD).

The gBOLD-CSF coupling strength showed a positive correlation with subjects’ age (*r* = 0.33, *p* = 0.01), i.e., the coupling strength was stronger (more negative) with decreasing age (Figure 2E). This gBOLD-CSF coupling relationship for our subjects aged between 24 to 68 years had the same pattern from the previous report with elderly subjects^[24, 27]^. However, there was no significant difference between males and females for the gBOLD-CSF coupling strength in the whole sample (*t*_(60)_ = 1.013, *p* = 0.315; Figure 2F). The gBOLD-CSF coupling strength of HCs was significantly stronger than the coupling of SCA3 patients (*t*_(60)_ = -2.08, *p* = 0.041; Figure 2A and Table 2), suggesting that glymphatic function deficiency may be involved in SCA3 patients.

**Table 2.** Comparisons of multiple glymphatic measurements between HCs and SCA3 patients.

| Glymphatic measurements | SCA3 | HCs | T values | p values |
| --- | --- | --- | --- | --- |
| **CSF volume** | N = 34 | N = 36 |  |  |
| CSF (ml) | 1.25±0.26 | 1.04±0.23 | 3.75 | <0.001 |
| CSF (ratio of ICV x 10<sup>3</sup>) | 0.83±0.14 | 0.69±0.12 | 4.44 | <0.001 |
| **CP volume** | N = 34 | N = 36 |  |  |
| CP (ml) | 1.19±0.4 | 0.91±0.47 | 2.64 | 0.01 |
| CP (ratio of ICV x 10<sup>3</sup>) | 0.78±0.23 | 0.6±0.28 | 2.87 | 0.006 |
| **BOLD-CSF coupling** | N = 27 | N = 35 |  |  |
| aBOLD-CSF | -0.22±0.11 | -0.27±0.13 | 1.54 | 0.128 |
| mBOLD-CSF | -0.23±0.11 | -0.34±0.18 | 3.06 | 0.003 |
| pBOLD-CSF | -0.23±0.12 | -0.28±0.16 | 1.41 | 0.165 |
| cbBOLD-CSF | -0.15±0.12 | -0.25±0.18 | 2.34 | 0.023 |
| gBOLD-CSF | -0.22±0.12 | -0.29±0.15 | 2.08 | 0.041 |
| **DTI-ALPS** | N = 28 | N = 35 |  |  |
| aDTI-ALPS | 1.18±0.13 | 1.26±0.18 | -2 | 0.05 |
| mDTI-ALPS | 1.24±0.2 | 1.39±0.18 | -3.1 | 0.003 |
| pDTI-ALPS | 1.13±0.16 | 1.19±0.18 | -1.41 | 0.165 |
| gDTI-ALPS | 1.16±0.12 | 1.27±0.15 | -3.02 | 0.004 |

Note: Data is presented as mean ± SD. HCs = Healthy controls, SCA3 = Spinocerebellar ataxia type 3, ICV = Intracranial volume, CSF = Cerebrospinal fluid, CP = Choroid plexus, a/m/p/cb/gBOLD-CSF coupling = The coupling between
anterior/middle/posterior/cerebellum/global cortical gray matter of blood-oxygen-level-dependent signals and cerebrospinal fluid signals; a/m/p/gDTI-ALPS = Anterior/middle/posterior/global diffusion tensor image analysis along the perivascular space.

### Group comparisons of regional glymphatic measurements between HCs and SCA3

To further confirm the glymphatic impairment in SCA3, multiple regional glymphatic differences were calculated between HCs and SCA3 patients. Between-group comparisons of regional glymphatic measurements (Table 2) showed that SCA3 patients had higher CP (*t*_(68)_ = 2.64, *p* = 0.01) and CSF (*t*_(68)_ = 3.75, *p* < 0.001) volumes than HCs, especially after controlling the inter-subject variability (*t*_(68)_ = 2.87, *p* = 0.006 for CP ratio and *t*_(68)_ = 4.44, *p* < 0.001 for CSF ratio). Additionally, when examining the regional DTI-ALPS between the two groups, mDTI-ALPS (*t*_(61)_ = -3.1, *p* = 0.003) and gDTI-ALPS (*t*_(61)_ = -3.02, *p* = 0.004) were statistically weaker in SCA3 patients than that in HCs. No significant difference was found in aDTI-ALPS (*t*_(61)_ = -2, *p* = 0.05) and pDTI-ALPS (*t*_(61)_ = -1.41, *p* = 0.165) between the two groups. This regional DTI-ALPS pattern was consistent with that of the regional BOLD-CSF coupling, which was also detected with significant differences in mBOLD-CSF coupling (*t*_(60)_ = 3.06, *p* = 0.003), cbBOLD-CSF coupling (*t*_(60)_ = 2.34, *p* = 0.023) and gBOLD-CSF coupling (*t*_(60)_ = 2.08, *p* = 0.041), but not in aBOLD-CSF (*t*_(60)_ = 1.54, *p* = 0.128) coupling and pBOLD-CSF coupling (*t*_(60)_ = 1.41, *p* = 0.165). These findings demonstrated that the glymphatic function in the midbrain, cerebellum, and middle regions was severely impaired in SCA3 patients.

### The spatial variation and correlation of regional glymphatic measurements

We then investigated how DTI-ALPS and BOLD-CSF coupling as indexes for glymphatic function varied spatially across cortical regions and fibers in HCs and SCA3 patients. In line with a prior report, DTI-ALPS gradients in both groups demonstrated an ascending trend from anterior to middle regions, followed by a decline from middle to posterior regions^[28]^ (Figure 3A). Significant differences were mainly found in the anterior (*t*_(61)_ range: 2.12 to 2.27, *p* range: 0.027 to 0.038) and middle regions (*t*_(61)_ range: 2.16 to 3.04, *p* range: 0.004 to 0.035) between HCs and SCA3 patients.

**Figure 3.**
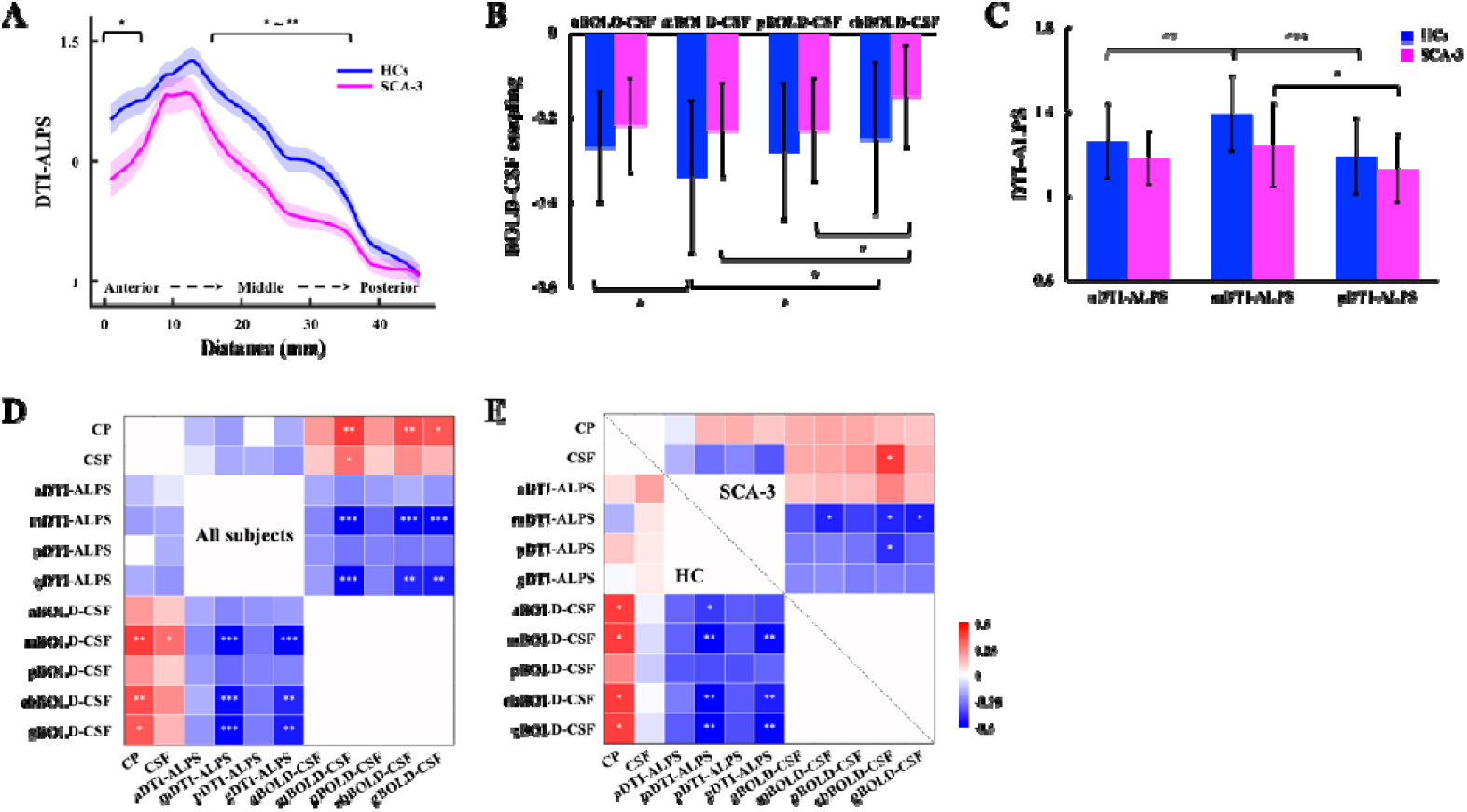
Glymphatic gradient and the relationship between different regional glymphatic function. (A) The dissociating gradient of DTI-ALPS from anterior to posterior regions between HCs (*N* = 35) and SCA3 patients (*N* = 28). Shaded areas are 95% interval of the glymphatic gradient within HCs and SCA3 patients. Blue and magenta lines denote HCs and SCA3 patients. (B) The distribution of BOLD-CSF coupling from anterior to cerebellum regions within HCs (*N* = 35) and SCA3 patients (*N* = 27). Blue and magenta bars denote HCs and SCA3 patients. Error bars represent the SD. (C) The distribution of DTI-ALPS from anterior to posterior regions within HCs (*N* = 35) and SCA3 patients (*N* = 28). Blue and magenta bars denote HCs and SCA3 patients. Error bars represent the SD. (D) and (E) The relationship among multiple glymphatic measurements in all subjects and within HCs and SCA3 patients, separately. *P* values in heatmaps were adjusted for age and sex. \**p* < 0.05, \*\**p* < 0.01, \*\*\**p* < 0.001.

In both groups, the mDTI-ALPS was stronger than the pDTI-ALPS (*t*_(68)_ = 4.65, *p* < 0.001 for HCs; *t*_(54)_ = 2.38, *p* = 0.021 for SCA3 patients; FDR correction), while the mDTI-ALPS surpassed the aDTI-ALPS only in HCs (*t*_(68)_ = 3.05, *p* = 0.003; FDR correction; Figure 3C). Moreover, the mBOLD-CSF coupling was stronger than the cbBOLD-CSF coupling in the two groups (*t*_(68)_ = -2.12, *p* = 0.038 for HCs; *t*_(52)_ = -2.25, *p* = 0.029 for SCA3 patients; FDR correction), with the mBOLD-CSF coupling being stronger than the aBOLD-CSF coupling exclusively in HCs (*t*_(68)_ = -2.1, *p* = 0.039; FDR correction), and the pBOLD-CSF coupling being stronger than cbBOLD-CSF coupling was observed solely in SCA3 patients (*t*_(52)_ = -2.24, *p* = 0.029 for SCA3 patients; FDR correction; Figure 3B). Furthermore, statistically significant positive correlations were primarily showed between mBOLD-CSF coupling, cbBOLD-CSF coupling, and gBOLD-CSF coupling with CP volume (*r* range: 0.33 to 0.39, *p* range: 0.013 to 0.042), along with negative correlations with mDTI-ALPS (*r* range: -0.53 to -0.48, all *p* <= 0.001) and gDTI-ALPS (*r* range: -0.49 to -0.43, *p* range: 0.001 to 0.009) across the entire subjects (Figure 3E and 3F). These results highlighted the different spatial pattern of glymphatic function between HCs and SCA3 patients.

### Relationship between aberrant glymphatic function and SCA3 symptoms

Our final analysis was to understand the association between altered glymphatic function and SCA3 symptoms. After adjusting for age and sex, and controlling for FDR, significant linear relationship was found between regional glymphatic measurements and SCA3 disease duration (*r* = 0.45, *p* = 0.03 for CP ratio; *r* = 0.6, *p* = 0.006 for gBOLD-CSF coupling; *r* = -0.56, *p* = 0.009 for gDTI-ALPS; Figure 4 A to D). Specifically, the CP ratio and CSF ratio were positively associated with ataxia and sleep assessments (*r* range: 0.37 to 0.53, *p* range: 0.002 to 0.033; Figure 4 A and B). All neuropsychological assessments for ataxia, psychiatry, and sleep symptoms were mainly significantly correlated with m/pBOLD-CSF (*r* range: 0.33 to 0.39, *p* range: 0.013 to 0.042) coupling strength, and mDTI-ALPS strength (*r* range: -0.65 to -0.33, *p* range: 0.001 to 0.047) with the exception of psychiatric assessments (Figure 4C and 4D), implying that the SCA3 disease severity increased with weakened glymphatic function. Further, these relationships indicated that impaired glymphatic function, as depicted by regional glymphatic measurements, could serve as predictive indicators for the clinical symptoms of SCA3.

**Figure 4.**
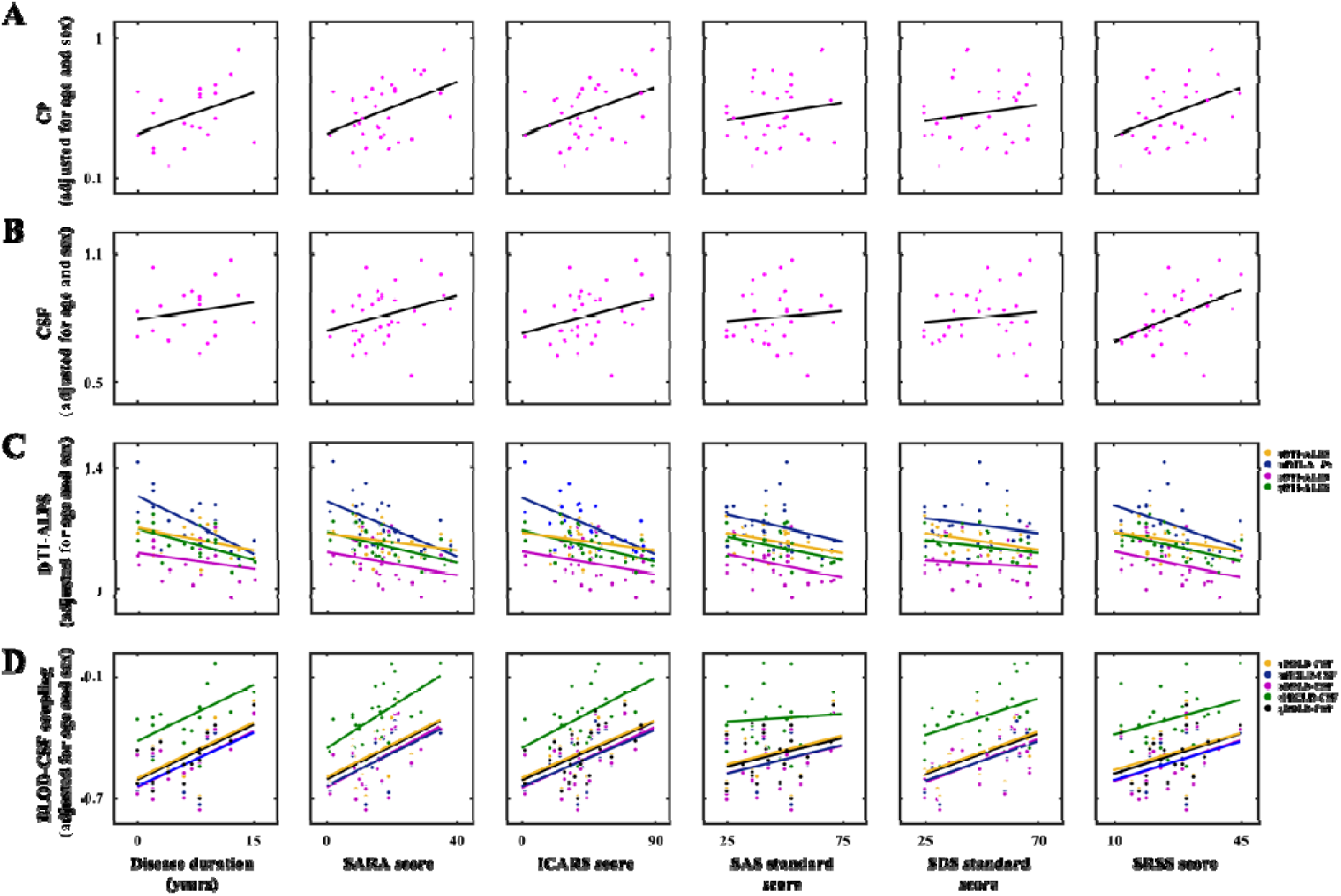
SCA3 symptoms were significantly correlated with different regional glymphatic function. (A-D) The CP ratio, CSF ratio, a/m/p/cb/gBOLD-CSF coupling and a/m/p/gDTI-ALPS adjusted for age and sex were correlated with different clinical assessments. Each magenta dot represents an SCA3 patient in the (A) and (B). Dark yellow, navy blue, deep purple, and dark green represent a/m/p/gDTI-ALPS in the (C). Dark yellow, navy blue, deep purple, dark green, and black represent a/m/p/cb/gBOLD-CSF coupling in the (D).

## Discussion

Drawing on the multiple glymphatic approaches, the current study aimed at comprehensively evaluating glymphatic function and its spatial distribution in patients with SCA3. These glymphatic approaches would enable us to estimate CSF morphology, CSF production, subcortical and cortical glymphatic movements using CSF volume, CP volume, DTI-ALPS, and BOLD-CSF coupling, respectively. Regional glymphatic function was then delineated to ascertain spatial patterns and correlated with clinical assessments in SCA3 patients. We demonstrated that glymphatic function was positively associated with age, mirroring the pattern reported previously in elderly cohort^[24, 27]^. Moreover, significant alterations in glymphatic function were observed, particularly within the midbrain, cerebellum, and middle regions, among SCA3 patients. These discernible deviations in regional glymphatic functions exhibited spatial variation, which were further associated with SCA3-related symptoms. Together, the current results suggested that the altered glymphatic functions affect pathophysiology of SCA3 in a spatially differentiated way, presumably through its effect on glymphatic clearance.

The present study showed abnormally elevated CSF and CP volumes in SCA3 patients compared to HCs. This enlargement in CSF and CP volume has been reported to be potentially linked to neuroinflammation and glymphatic impairment in neurodegenerative diseases^[28, 31, 32]^ and psychosis spectrum^[45, 46]^. Therefore, the impaired glymphatic clearance mechanisms in SCA3 may mitigate glymphatic dysfunction by enhancing waste clearance through increased production and transmission of functional units in the CSF and CP^[47]^. Furthermore, the between-group comparison results demonstrated that in contrast to HCs, SCA3 patients manifested weaker m/cb/gBOLD-CSF coupling and lower m/gDTI-ALPS. In line with the previous studies, SCA3 patients in the current study exhibiting predominant atrophic changes in the precentral gyrus, paracentral lobes, and cerebellum also witnessed the deposition of toxic proteins (e.g., tau and polyglutamine protein)^[6, 17, 18, 21]^ and neuro-damage markers^[19, 20]^. The bidirectional flow within the glymphatic system may facilitate the clearance of aberrant proteins from the ISF into the ventricular or subarachnoid compartments^[48, 49]^. Consequently, slowed glymphatic movement can result in waste accumulation across brain cortex and subcortex in SCA3 patients. Additionally, these findings also revealed impairment of glymphatic function within midbrain, cerebellum, and middle regions in patients with SCA3. The pathological underpinnings of SCA3 were primarily localized to the cerebellum-neostriatum-motor and association cortical circuits^[50]^, consistent with the distributional pattern of glymphatic disfunction.

Another noteworthy finding from the current research was that, in regional glymphatic distributions we tested, the cortical and subcortical clearance showed an ascending trend from anterior to middle regions, followed by a descending trend from middle to posterior and cerebellum regions. The spatial dynamics in subcortical and cortical clearance were consistent with the anterior-posterior gradient observed in white matter microstructural diffusivity^[51]^ and the motor-sensory dominant pattern of regional BOLD^[25, 52]^, respectively. Given the consistent utilization of the same CSF signal for each subject, any spatial variation in the regional BOLD-CSF coupling can be exclusively ascribed to alterations in regional BOLD signals. This pattern of brain co-activation was concomitant with specific deactivation in subcortical regions associated with arousal regulation, notably the basal ganglia and brainstem^[25, 52]^, and was strongly related to SCA3 etiology since the accumulation and aggregation of expanded polyglutamine stretches within susceptible brain nuclei may precipitate direct or indirect neurotoxic effects, ultimately culminating in neuronal loss and cerebral atrophy^[6, 17, 18]^. Furthermore, in mice, neural response in the sensory motor cortex induced by whisker stimulation was recently found to be accompanied by accelerated CSF flow in PVS^[53]^. This suggested that neurovascular coupling was involved in both supplying metabolites and removal of their waste products. Therefore, inadequate regional glymphatic clearance may lead to the accumulation and dissemination of neurotoxic proteins within the he midbrain, cerebellum, and middle regions, thereby exacerbating the pathogenesis of SCA3.

With regard to the relationship between multiple glymphatic approaches and SCA3 symptoms, our results demonstrated significant correlations between these glymphatic indexes and SCA3-related symptoms as well as the disease severity. These findings highlighted the potential of glymphatic dysfunction as a biomarker for disease severity and progression of SCA3. Similar associations have been reported in other neurodegenerative diseases, emphasizing the pervasive impact of glymphatic dysfunction on neurological function^[24–28, 31–36]^. Furthermore, the resting-state BOLD signal performed sleep dependence similar to glymphatic function, which was notably strong during drowsiness and sleep states^[54, 55]^. Thus, the sleep disturbances in SCA3 patients may cause impaired glymphatic clearance, resulting in the weaker glymphatic indexes. Further investigations could extend this line of research to explore whether enhanced glymphatic clearance could improve SCA3 symptoms.

Finally, some caveats need to be noted regarding the present study. Our study did not directly evaluate the accumulation of neurotoxic wastes (proteins or small molecules). Further studies adopting PET data of the relevant proteins (e.g., tau-PET images) could provide empirical evidence with the relationship between regional glymphatic function and proteins deposition, especially in the midbrain, cerebellum, and middle regions. Moreover, as subjects in the present study consisted of a cohort, we were unable to offer insights into the longitudinal changes of spatial distribution of the glymphatic function in SCA3. As such, our data warrants future follow-up studies, with larger sample sizes and in independent cohorts, to validate and extend these findings.

In summary, drawing on multiple glymphatic measurements, the current study provided comprehensive and compelling evidence for the involvement of glymphatic dysfunction in SCA3 patients. Notably, SCA3 patients exhibited spatial specificity of glymphatic dysfunction in midbrain, cerebellum, and middle regions, which revealed the pathological patterns of SCA3. Furthermore, the regional glymphatic alterations were closely associated with the SCA3 symptoms. Our findings provided valuable insights into the interplay between glymphatic dysfunction, brain structural alterations, and clinical symptoms, contributing to a deeper understanding of the pathophysiology of SCA3.

## Supporting information

SupportingInformation

## Ethics statement

This study was conducted in accordance with the Declaration of Helsinki (as revised in 2013) and was approved by the Ethics Committee of the China-Japan Friendship Hospital. All the participants and/or their relatives were informed about this study and provided their written informed consent. The information of all participants had been fully anonymized.

## Acknowledgements

This work was supported by Macao Science and Technology Development Fund (No. 0020/2019/AMJ and 0011/2018/A1), the University of Macau (No. MYRG2020-00067-FHS, MYRG2019-00082-FHS, and MYRG2018-00081-FHS), Higher Education Fund of Macao SAR Government (No. CP-UMAC-2020-01), National Natural Science Foundation of China (No. 82271953 and 81971585), Guangzhou Science and Technology Planning Project (No. 202103010001), STI2030-Major Projects (No. 2022ZD0213300), Capital’s Funds for Health Improvement and Research (No. 2022-1-2031) and Beijing Municipal Science and Technology Project (No. Z211100003521009) and Open Research Fund of the State Key Laboratory of Cognitive Neuroscience and Learning.

## Author contribution

L.H.: conceptualization, methodology, validation, investigation, formal analysis, writing—original draft, writing—review and editing. MX.X: conceptualization, methodology, validation, investigation, formal analysis, writing—original draft, writing—review and editing. LW.Z.: methodology, resources, data curation, validation. F.G.: methodology, validation, writing—review and editing. LX.Z.: methodology, validation, formal analysis. AC.Y.: methodology, data curation. JX.L.: methodology, data curation. A.S.: writing—review and editing. GL.M.: conceptualization resources, funding acquisition, project administration, writing—review and editing. Z.Y.: conceptualization resources, funding acquisition, project administration, writing—review and editing.

## Supplementary material

Supplementary material is available in supplementary files.

## Data availability

The data that support the findings of this study are available from the corresponding authors upon reasonable request.

## Code availability

All analyses used open-source software with URL links already included in Methods. Code used in the analyses described in this paper will be made available upon acceptance of the manuscript.

