## SupportingInformation for "Regional Glymphatic Dysfunction is linked to Spinocerebellar Ataxia Type 3 pathophysiology"

**Supporting Information**

**Subjects**

From June 2021 to April 2024, a prospective cohort of 71 patients with spinocerebellar degeneration patients were recruited from the outpatient clinic of the China-Japan Friendship Hospital. Genetic testing was conducted to exclude those diagnosed with Multiple System Atrophy (MSA), Autosomal Recessive Cerebellar Ataxia (ARCA), and other types of SCA. 34 symptomatic patients with genetically confirmed SCA3 were included in the analysis. All patients underwent integrated neuropsychological tests and MRI examinations. The Scale for the Assessment and Rating of Ataxia (SARA) and the International Cooperative Ataxia Rating Scale (ICARS) scores were examined and recorded by trained neurologists to measure the disease severity of SCA3 patients^[1]^. Additionally, healthy controls (HCs) matched for age, gender, and educational level were recruited from the local community.

The inclusion criteria of SCA3: (1) diagnosed as SCA3 by expert neurologists according to the family history, neurological manifestations, genetic and molecular tests, and routine brain MRI findings; (2) complete clinical and imaging data; (3) age≥18 years; (4) right-handed. The exclusion criteria were as follows: (1) challenging to cooperate during MRI examination, and the image quality is too poor for image analysis; (2) a history of other brains organic and metabolic diseases; (3) pregnant and lactating women; and (4) other MRI contraindications.

**Quality Control**

To ensure the quality of imaging data, T1-weighted sMRI, rs-fMRI and DTI data exhibiting excessive motion, artifacts or poor brain coverage were excluded. Moreover, rigorous quality control (QC) was performed for individual sMRI scans following CAT12 preprocessing. Several parameter-specific ratings were provided as well as a handy overall rating for quantifying image qualities such as noise, intensity inhomogeneities, image resolution, and weighted average (IQR) ^[2]^. Subjects with poor MRI quality, as indicated by any of the QC parameters being lower than B, were also excluded. For the rs-fMRI data, outlier detection was performed for each subject, in which framewise displacement (FD) and root-mean-square of voxel-wise differentiated signal (DVARS) were estimated using fsl_motion_outliers implemented in the FSL software (version 6.0.5; <https://fsl.fmrib.ox.ac.uk/fsl/fslwiki>). FD was derived as the cumulative sum of the absolute values of all six translational and rotational realignment parameters^[3]^. Simultaneously, DVARS was computed as the root mean square (RMS) value, calculated at each volume, representing the first derivatives of all voxel time series across the entire brain^[3, 4]^. Volumes with mean FD values exceeding 0.3 mm or DVARS values surpassing 50 were marked as outliers (censored frames). One frame before and two frames after these outliner volumes were also flagged as censored frames, together with those lasting fewer than five contiguous volumes. Subjects who had more than half the volumes labeled as censored frames were rejected from further analysis. In addition to these automated quality control measures, all preprocessed rs-fMRI time series underwent visual inspection for artifacts, especially motion artifacts that might affect the bottom slice of rs-fMRI data. DTI quantitative identification of slices with signal loss was performed using the eddy implemented in the FSL software (version 6.0.5; <https://fsl.fmrib.ox.ac.uk/fsl/fslwiki>). Outlier volumes were replaced by non-parametric predictions using the Gaussian process^[5]^. Finally, 36 HCs and 34 SCA3 patients with structural MRI were used to obtain the CP and CSF volumes. 1 HC and 7 SCA3 patients with rs-fMRI as well as 1 HC and 6 SCA3 patients with DTI were excluded for further BOLD-CSF coupling and DTI-ALPS analysis.

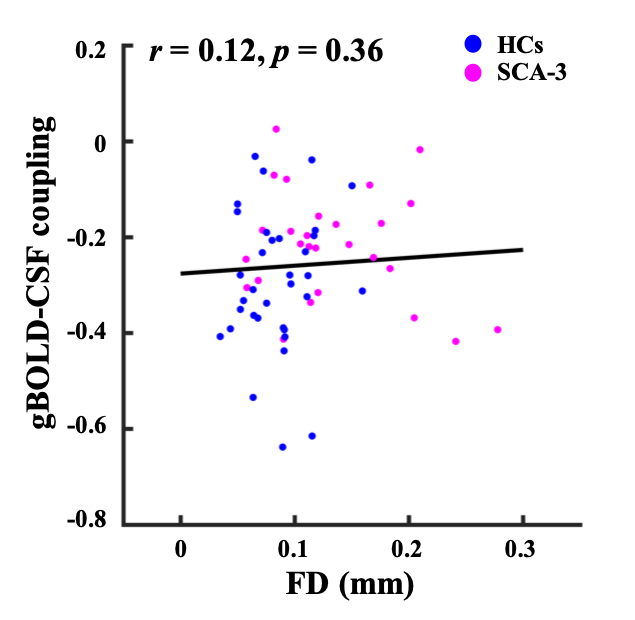

**Figure S1. The head motion did not affect the BOLD-CSF coupling strength.** The head motion of subjects, quantified by mean FD, was not correlated with the BOLD-CSF coupling strength (*r* = 0.12, *p* = 0.36).
